# Single Cell Biological Microlasers Powered by Deep Learning

**DOI:** 10.1101/2021.01.21.427584

**Authors:** Zhen Qiao, Wen Sun, Na Zhang, Randall Ang Jie, Sing Yian Chew, Yu-Cheng Chen

## Abstract

Cellular lasers are cutting-edge technologies for biomedical applications. Due to the enhanced interactions between light and cells in microcavities, cellular properties and subtle changes of cells can be significantly reflected by the laser emission characteristics. In particular, transverse laser modes from single-cell lasers which utilize Fabry–Pérot cavities are highly correlated to the spatial biophysical properties of cells. However, the high chaotic and complex variation of laser modes limits their practical applications for cell detections. Deep learning technique has demonstrated its powerful capability in solving complex imaging problems, which is expected to be applied for cell detections based on laser mode imaging. In this study, deep learning technique was applied to analyze laser modes generated from single-cell lasers, in which a correlation between laser modes and physical properties of cells was built. As a proof-of-concept, we demonstrated the predictions of cell sizes using deep learning based on laser mode imaging. In the first part, bioinspired cell models were fabricated to systematically study how cell sizes affect the characteristics of laser modes. By training a convolutional neuron network (CNN) model with laser mode images, predictions of cell model diameters with a sub-wavelength accuracy were achieved. In the second part, deep learning was employed to study laser modes generated from biological cells. By training a CNN model with laser mode images acquired from astrocyte cells, predictions of cell sizes with a sub-wavelength accuracy were also achieved. The results show the great potential of laser mode imaging integrated with deep learning for cell analysis and biophysical studies.

## 1. Introduction

Biological laser, which embeds biological materials into the gain medium and optical cavity, is an emerging technology that offers new ways of using laser light in biomedical applications^1–18^. Thanks to the strong light-matter interactions provided by an optical cavity, subtle changes in biological gain media could be amplified, leading to distinctive output lasing characteristics. Recent advances in cellular lasers have attracted tremendous interest in cell tracking, intracellular detection, and phenotyping by exploiting various types of cavities^5–9^. In particular, single-cell lasers that utilize Fabry-Pérot (FP) microcavities possess the advantages of whole-body interaction between the electromagnetic field and the gain medium, providing a sensitive approach to study the physical structures and interactions in cells^4, 5, 19^. One of the most peculiar features of such lasers is the “lens effect” of cells introduced in FP cavities, in which the laser patterns are formed by the superposition of a series of transverse laser modes^20^. While most studies mainly focused on the spectral information of biological lasers, less attention was paid to the spatial information of transverse laser modes. In principle, the attributes of laser modes (e.g., mode types and orders) are highly correlated to cells' biophysical properties, for instance, size, refractive index distribution, cell morphology, scattering, and fluorophore (gain) distribution in the cell. As typical transverse modes, Hermite Gaussian (HG), Laguerre Gaussian (LG) and Ince Gaussian (LG) modes are mostly observed in single cell lasers^4, 20^. The different patterns of laser modes can be viewed as integrated information of the single cell confined in the cavity. As such, laser mode imaging of single cells possesses a huge potential for studying cellular properties and changes in cells. However, the high chaos and huge complexity of laser modes remain the key challenge toward practical applications.

In recent years, deep learning technology has demonstrated its powerful capability in a wide range of complex data analysis, particularly in bio-imaging^21–27^. By learning intrinsic features of input images using conventional neural networks (CNN), a multi-layer mapping function can be built between measured information and input imaging data. Such end-to-end frameworks can deliver high-accuracy predictions of the underlying information in images^28, 29^, thereby solving imaging problems associated with complicated physical processes in biology and medicine. Inspired by deep learning analysis, it is envisaged that laser mode images generated from single cell lasers could be trained with CNN and analyzed more precisely for single cellular studies. Through deep learning analysis, the hidden information in laser modes may be revealed.

In this study, deep learning technology was applied to reveal the complex laser modes generated from cells sandwiched in a FP cavity, in which the physical properties of bioinspired cell models and biological cells were both investigated. As a proof-of-concept, we demonstrated the possibility of employing deep learning method to predict the sizes of cells based on laser mode imaging. In the first part, bioinspired (artificial) cells were prepared to systematically study the fundamental characteristics of laser modes. The diameters of cell models were subsequently predicted from the corresponding laser mode images using CNN models. Consequently, predictions of cell model diameters with a sub-wavelength accuracy were achieved. In the second part, a similar method was employed to study laser modes generated from astrocyte cells derived from mice nervous system. By training the deep learning model with laser mode images acquired from astrocyte cells, predictions of cell sizes with a sub-wavelength accuracy were also achieved. The results reveal the great potential of laser mode imaging, offering new possibilities for applications in single cell analysis and biophysical studies.

## 2. Results

### 2.1 Concept of laser mode imaging of cells powered by deep learning

Figure 1 illustrates the generation of laser modes from single cell lasers and the schematic of laser mode imaging of single cells powered by deep learning. The single cell lasers were fabricated using Fabry-Pérot (FP) microcavities, in which a dye-doped bioinspired cell model or a biological cell was sandwiched by two cavity mirrors (Fig. 1a). Due to lens effect induced by cells in FP cavities, transverse laser modes will be generated from single cells when pumped at the extinction wavelength corresponding to its gain medium (fluorophore). In order to resolve the output laser modes, hyperspectral images of laser modes are measured using an imaging spectrometer (Fig. 1b), in which the laser mode patterns with different lasing wavelengths can be obtained. Figures 1c and 1d show examples of laser emissions acquired from bioinspired cell models and biological cells, respectively. The far-field laser emission patterns (left in Figs. 1c and 1d) represent the superposition of a series of transvers laser modes with different lasing wavelengths. The hyperspectral images of laser modes show complex laser emission patterns with various laser mode components (right in Figs. 1c and 1d). The lasing spectra (Figs. 1c and 1d) presents the corresponding lasing wavelengths of the transverse laser modes. To further demonstrate the lasing operations, spectrally integrated output intensities as a function of pump energy density were also measured (insets in Figs. 1c and 1d), which presents clear threshold behaviors of single cell lasers. Due to the huge complexity of laser modes, deep learning technique was applied to establish the correlation between various laser modes and biological cells. Explicitly, the laser mode images extracted from large number of cells were used to train a convolutional neural network (CNN) model; subsequently, the physical properties of cells were predicted by the trained CNN model (Fig. 1e). As a proof of concept, we demonstrated the possibility of employing deep learning method to predict the cell sizes with a sub-wavelength accuracy.

**Figure 1.**
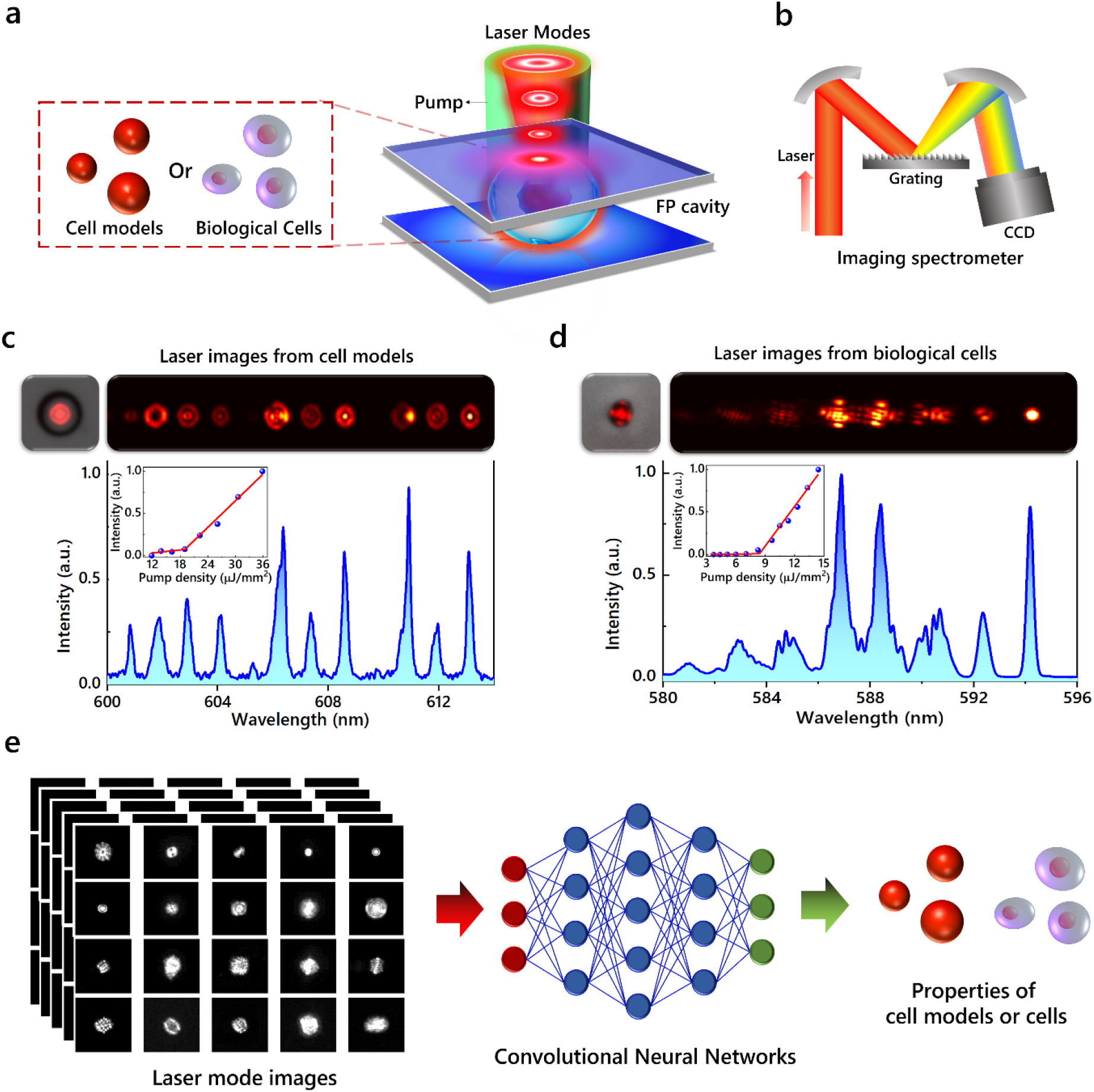
Schematic illustration of laser mode imaging of cells powered by deep learning. **(a)** Schematic of single cell laser that utilizes Fabry–Pérot (FP) microcavity. A dye-doped bioinspired cell model or a biological cell was sandwiched within a FP cavity. Laser modes emit from the FP cavity after pumping. **(b)** Schematic of hyperspectral imaging setup. Hyperspectral images of laser modes were obtained using an imaging spectrometer, in which a CCD was used to record the laser modes with different wavelengths after the laser emission beam was diffracted by a grating. **(c) (d)** Laser modes generated from bioinspired cell models (c) and biological cells (d). Left: far-field laser emission patterns which are superimposed on the bright-field images of bioinspired cell models (c) and biological cells (d). Right: hyperspectral images of laser modes and the corresponding lasing spectra. Insets in (c) and (d): spectrally integrated output intensities as a function of pump energy density. **(e)** Schematic of laser mode imaging of cells powered by deep learning. The laser mode images extracted from (c) and (d) were used to train a CNN model; consequently, physical properties of bioinspired cell models and biological cells were predicted based on the corresponding laser mode images using the trained CNN model.

### 2.2 Laser modes generation from bioinspired cell models

Firstly, bioinspired cell models were used to systematically investigate how the sizes of cell models affect the emitted laser modes from FP lasers. The cell models were fabricated by cholesteric liquid crystals (CLC) droplets doped with 4-(dicyanomethylene)-2-methyl-6-(4-dimethylaminostyryl)-4H-pyran (DCM) dye. Figure 2a shows the bright-field image and fluorescent image of a cell model (CLC droplet). As depicted in Fig. 2b, the cell models were sandwiched within a FP cavity, in which glass microspheres were used as spacers to fix the cavity length (Fig. 2b). By pumping at 480-nm wavelength with a fixed pump energy density, we recorded the laser modes generated from CLC droplets with different diameters. For instance, Fig. 2c shows the output laser modes from droplets with three different diameters (12.72 μm, 13.97 μm and 15.3 μm). The far-field laser emission patterns (left part in Fig. 2c) cover larger areas and possess more complex spatial structures for larger cell models. Meanwhile, the hyperspectral images (right part in Fig. 2c) show the corresponding laser mode components of the laser emission patterns. These laser modes possess centro-symmetrical intensity distributions, which can be well described by LG_*pl*_ modes, where *p* and *l* are the radial and azimuthal orders of LG modes, respectively. For instance, only LG_00_ and LG01 modes were obtained for a droplet with diameter of 12.72 μm. For larger droplets with a diameter of 13.97 μm, the FP laser emitted LG_00_, LG_01_, LG_10_ and LG_11_ modes. For an even larger droplet with diameter of 15.30 μm, much more laser modes with higher orders were generated, including LG_00_, LG01, LG_10_, LG_11_, LG_13_, LG_20_, LG_21_ and LG_30_ modes. Figure 2c presents the tendency that more laser modes with higher orders can be excited with the increasing cell model sizes.

**Figure 2.**
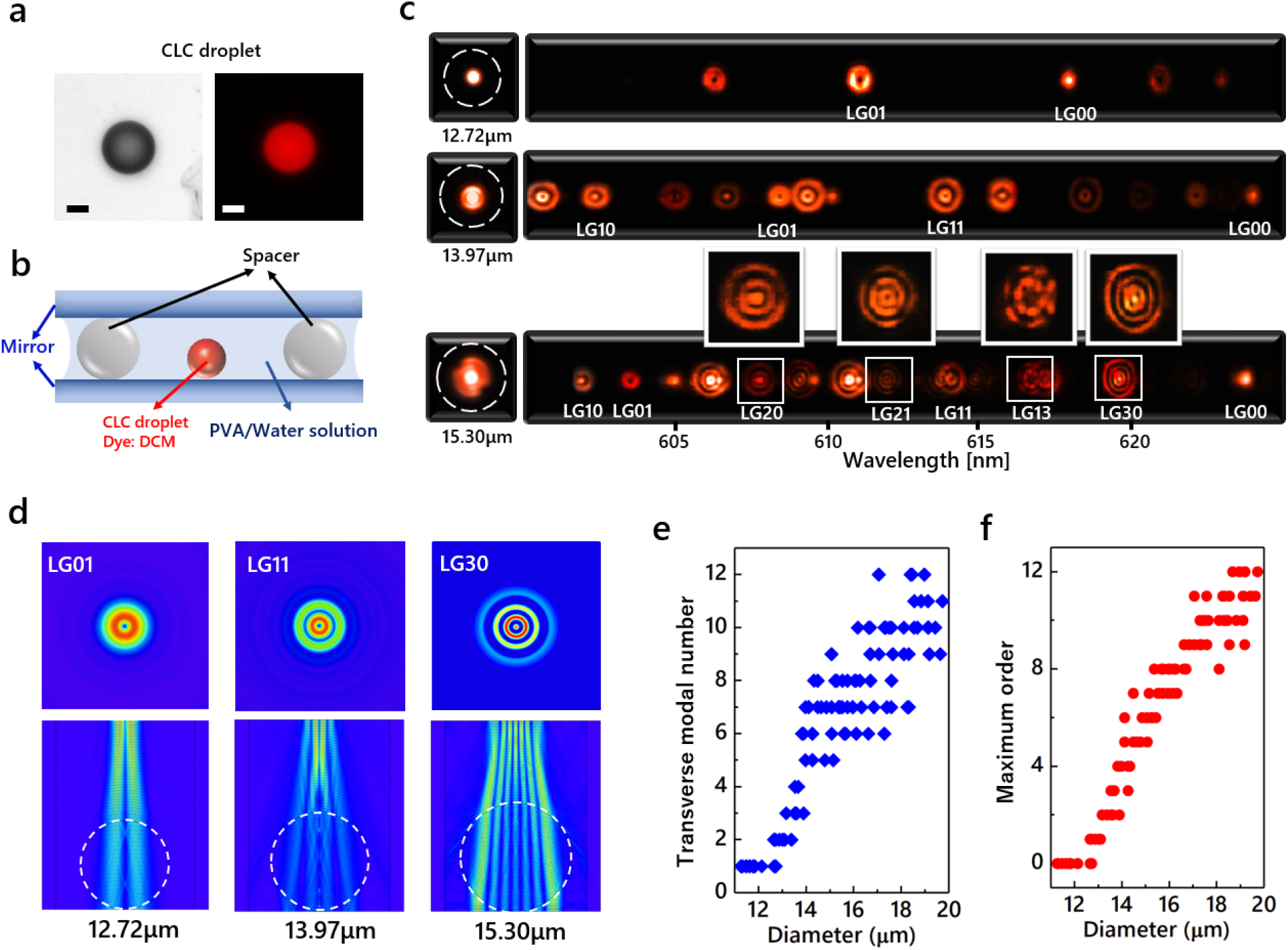
Laser mode generation from bioinspired cell models. **(a)** Bright-field image (left) and fluorescent image (right) of a dye-doped cell model. The cell models were fabricated by spherical cholesteric liquid crystal (CLC) droplets doped with DCM dye. Scale bar: 10 μm. **(b)** Schematic of a FP cavity sandwiching a DCM-doped CLC droplet. The surrounding environment is PVA/water solution. Microspheres with 27-μm diameters were used as spacers to fix the cavity length. **(c)** Far-field laser emission patterns (left) and the corresponding hyperspectral images of laser modes (right) generated from cell models with diameters of 12.72 μm, 13.97 μm and 15.30 μm. The generated laser modes are LG modes denoted by LG_*pl*_, where *p* and *l* are the orders of LG modes. White dashed circles: cell model outlines. **(d)**Simulations of laser mode generation from FP cavities sandwiching droplets with diameters of 12.72 μm, 13.97 μm and 15.30 μm. White dashed circles: cell model outlines. Top: transverse intensity distributions of LG_01_, LG_11_ and LG_30_ modes. Below: Side-view electric field distributions of LG_01_, LG_11_ and LG_30_ modes in FP cavities. **(e)** Measured transverse modal number of laser modes as a function of cell model diameter. For example, LG_00_ and LG_01_ modes were excited from a 12.72-μm diameter CLC droplet in (c); thus, the number of laser mode type is 2. **(f)** Measured max order of laser modes (*N*) as a function of cell model diameter. For LG_*pl*_ modes, *N*=2*p*+*l*; for IG_*pm*_ modes, *N*=*p*.

To understand the experimental results in Fig. 2c, theoretical simulations were performed on laser modes generation under different diameters. Three spherical droplets whose diameters are the same as the experiment (Fig. 2c) were considered in FP cavities. By numerically solving Helmholtz equation that governs laser oscillations in cavity, eigen solutions of transverse laser modes with maximum orders were obtained. As shown in Fig. 2d, the maximum order of laser modes that can form resonance in a FP cavity were found to be LG_01_, LG_11_ and LG_30_ modes for a droplet with diameter of 12.72, 13.97, and 15.3 μm, respectively. The simulation results showed good agreement with the experiment in Fig. 2c. As such, more laser modes with higher orders can be excited from larger cell models.

We further performed the statistics of the transverse modal numbers and maximum order of laser modes generated from cell models. It should be mentioned that, in addition to LG modes shown in Fig. 2c, IG modes can also be excited and should be considered. For unification, we define the order of laser modes by one index *N* which equals 2*p+l* for LG_*pl*_ modes; and *p* for IG_*pm*_ modes. For instance, LG_00_ and LG_01_ modes were both generated from a droplet with the diameter of 12.72 μm (Fig. 2c), in which LG_01_ mode is the maximum order of laser mode; therefore, the transverse modal number is 2 and the max order of the generated laser mode is 1 (N=1). Based on this method, the statistic results are summarized in Figs. 2e and 2f. Both the transverse modal number and the maximum order of laser modes presented an increasing tendency with the size of the droplets sandwiched in FP cavity. This result indicates that the maximum order and patterns of laser modes is highly dependent on the size of the target (droplet) inside the cavity.

### 2.3 Diversity of laser modes from bioinspired cell models with the same diameter

In principle, laser modes are extremely sensitive to spatial physical changes inside an optical cavity. In addition to the size (diameter) of cells, other spatial characteristics can also influence the emitted laser modes, such as refractive index distributions, structure-induced loss distributions, and gain (dye) distributions. Here we show the diversity and complexity of output laser modes based on different droplets with predefined size (diameter). Figure 3a depicts far-field laser emission patterns from six individual cell models with the exact same diameter (14.84μm).

**Figure 3.**
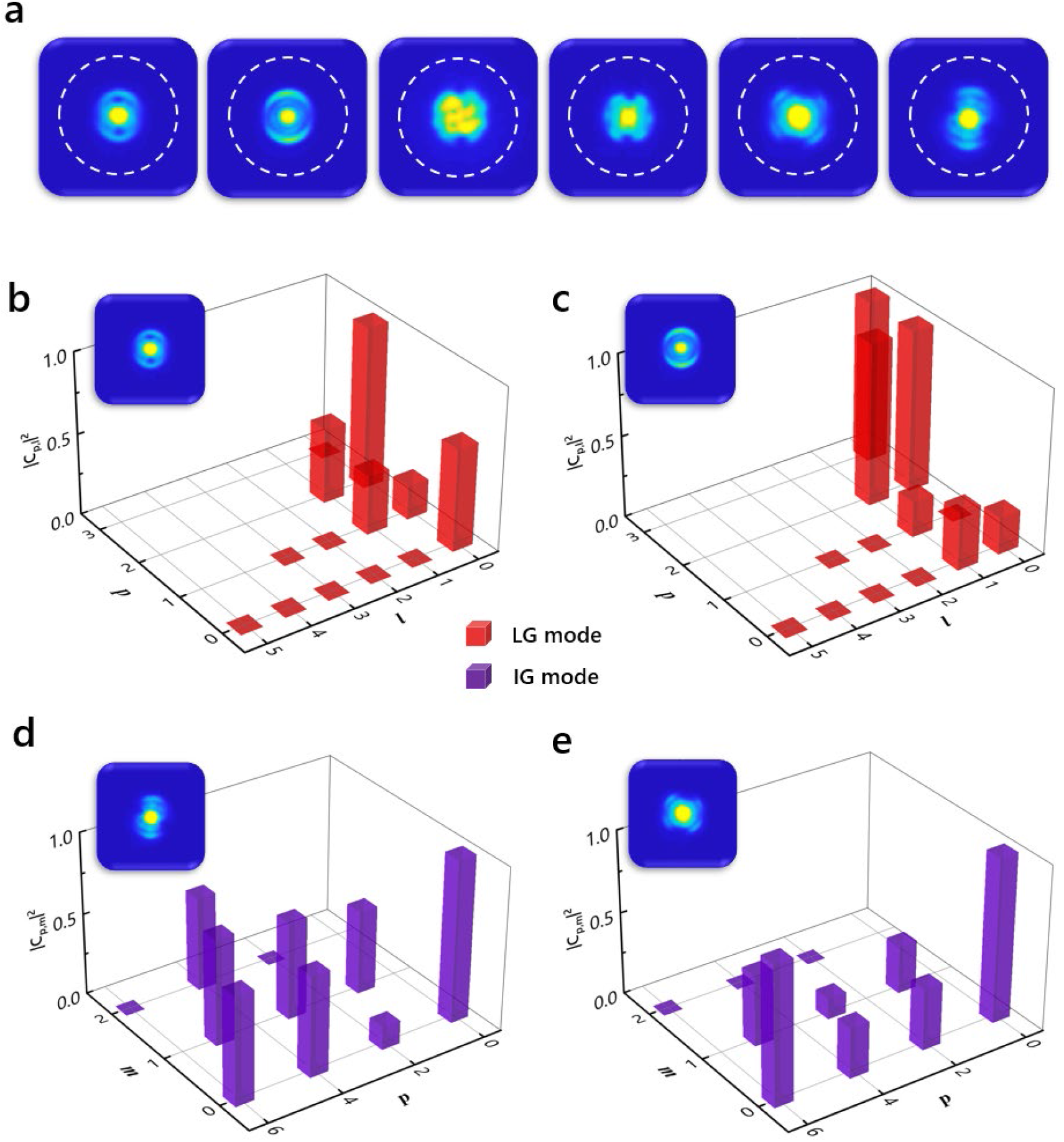
Diversity of laser modes from bioinspired cell models with the same diameter. **(a)** Examples of far-field laser emission patterns from the cell models (CLC droplets) with the same diameter of 14.84 μm. White dashed circles: cell model outlines. **(b) (c) (d) and (e)** Measured modal spectra of the laser emission patterns from four cell models with the same diameter of 14.84 μm. Insets in (b)~(e): laser emission patterns. |*c*_*p,l*_|^2^ is the normalized modal power of LG_*pl*_ mode; |*c*_*p,m*_|^2^ is the normalized modal power of IG_*pm*_ mode.

Apparently, the laser emission patterns derived from different droplets possess distinctive spatial intensity distributions. To further show the diversity of laser modes, transverse modal spectra of laser emissions were also measured in Figs. 3b-3e based on the hyperspectral images of laser modes. The laser emission patterns were determined by the intensities of the superposed laser mode components. By comparing Figs. 3b and 3c, the two FP lasers with same droplet size emitted LG modes; however, the fluctuated intensities of laser mode components leaded to different laser emission patterns (insets of Figs. 3b and 3c). Another set of comparison is shown in Figs. 3d and 3e, where the different intensity distributions of IG modes resulted in different laser emission patterns (insets of Figs. 3d and 3e). Such diversity of output laser modes makes it a challenge to retrieve or identify the critical information from laser modes. However, it is envisaged that laser modes could potentially become a powerful tool for cell analysis with the assistance of deep learning.

### 2.4 Size prediction of bioinspired cell models from laser mode imaging enabled by deep learning

In this section, we demonstrated the possibility of using deep learning method to predict the diameters of cell models based on laser mode imaging. To establish the accurate correlation between laser mode images and cell model diameters, laser modes from different sizes of droplets (cell models) were systematically analyzed. More than 2000 laser mode images were collected from droplets whose diameters range from 11~27μm. Considering that a FP laser emits a series of transverse laser modes, the spatial intensity distributions of laser emission patterns can be described as the following equation:

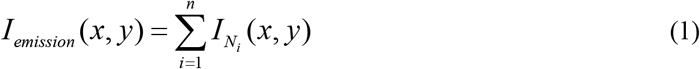

Here (*x*,*y*) demotes a spatial position; *I*_emission_ represents the spatial intensity distribution of a laser emission pattern; *I*_*Ni*_ is the spatial intensity distribution of the *i*-th laser mode component, where *N*_*i*_ denotes the order of *i*-th laser mode component; and *n* is the transverse modal number. As demonstrated in Figs. 2e and 2f, two parameters in Eq. 1 are highly correlated to the diameter of cell models: one is the transverse modal number (*n*), the other is the maximum order of laser modes [*N*_max_=Max (*N*_1_, *N*_2_, ..., *N*_*n*_)]. Therefore, the far-field laser emission pattern (formed by the superposition of *n* laser modes) and the corresponding max-order laser mode component, were selected as laser mode images to train the CNN model (Fig. 4a). Consequently, the input laser mode images possess the following intensity distributions:

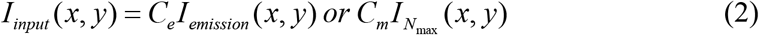

where *I*_*input*_ is the spatial intensity distributions of laser mode images (right in Fig. 4a) and *C*_*e*_, *C*_*m*_ are constants.

**Figure 4.**
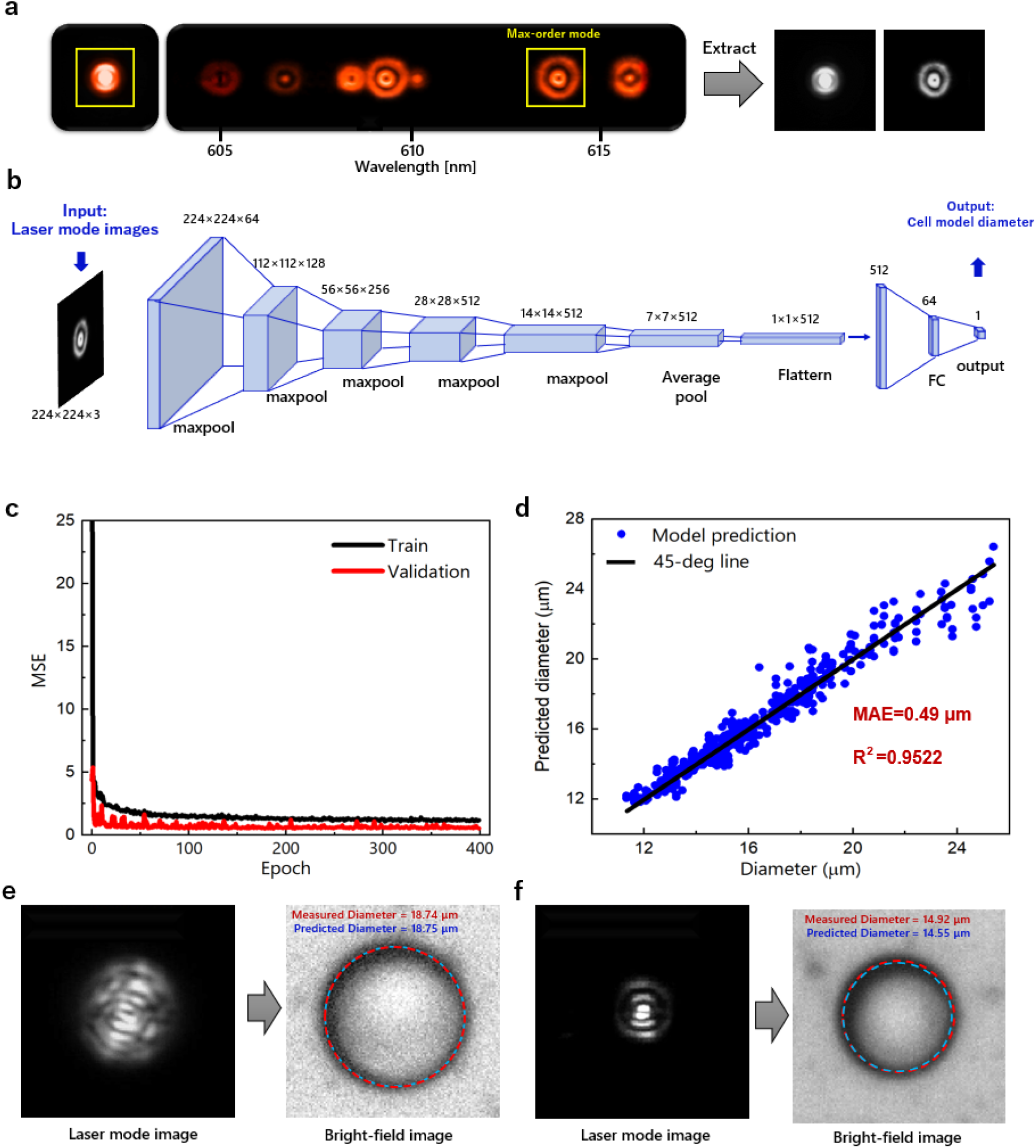
Size prediction of bioinspired cell models based on laser mode images using deep learning. **(a)** Selections of laser mode images. The far-filed laser emission patterns and the corresponding max-order laser mode components were extracted as laser mode images for training a CNN model. Yellow boxes: Selected laser mode images. **(b)** The CNN architecture based on VGG19 network. The boxes indicate the number and size of extracted feature maps from hidden layers. Input: laser mode images. Output: diameters of cell models. **(c)** The mean square error (MSE) for training and validation sets with respect to the epochs. **(d)** Scatter plot showing test set predictions made by the trained CNN model. Blue dots: predictions of cell model diameters. Black line: 45-degree line which represents the measured cell model diameters. The prediction mean absolute error (MAE) is 0.49 μm; the prediction R^2^ value is 0.9522. **(e) (f)**Two examples of laser mode images (left) and the corresponding predicted cell model diameters (right). Red dashed circles in bright-field images: measured outlines of cell models. Blue dashed circles in bright-field images: reconstructed outline of cell models based on the predicted diameters.

By training the CCN model with extracted laser mode images, the diameters of cell models (droplets) were predicted based on laser mode imaging. The deep learning network structure is shown in Fig. 4b, which was designed based on VGG19 (details in Method). In particular, the CNN was trained with 1567 laser mode images in the training dataset and validated with 223 images in the validation dataset. The mean squared errors (MSE) between measured and output diameters for the training and validation datasets with respect to the epochs are plotted in Fig. 4c. Both curves present convergence to low values with increasing number of epochs. To show the performance of the CNN model, 447 laser mode images were used to test the prediction performance. The scatter plots in Fig. 4d shows the test dataset predictions conducted by the trained CNN model with respect to the measured diameters. It can be observed that the predicted points are well distributed along the 45-degree line which shows the equality between measured and predicted diameters. We further calculated the mean average error (MAE) and *R*^2^ value based on the prediction data (details in Method), which are 0.49 μm and 0.9522, respectively. The small MAE demonstrated a sub-wavelength prediction accuracy made by the CNN model. In addition, the *R*^2^ value is close to 1, which further indicates a good regression result. Two examples from the test dataset are presented in Figs. 4e and 4f. The left panel represents the input laser mode images; the right panel shows the comparison between predicted and measured droplet diameters. The reconstructed profiles based on the predicted diameter were found to be very close to the measured profiles of cell models, showing excellent predictions made by the CNN model.

### 2.5 Laser modes generation from biological cells

Next, we explored the feasibility of employing deep learning for laser mode imaging of real biological cells. Figure 5 shows the laser mode generation from single cells by using astrocyte cells extracted from nervous systems of mice pups (Fig. 5a). All the cultured astrocyte cells were suspended and fixed before staining with Rhodamine 6G (Rh6G) (see Methods for details). The fluorescence microscopic images of single astrocyte cells are presented in Fig. 5a. Different from the spherical cell models, most astrocyte cells possess irregular shapes. Single cell micro-lasers were fabricated by sandwiching the astrocyte cells within FP cavities, in which microspheres were used as spacers to fix the cavity length. Note that Rh6G dye has a strong binding affinity to mitochondria in cells; however excessive dye molecules will also exist in the surrounding medium (Fig. 5b).

**Figure 5.**
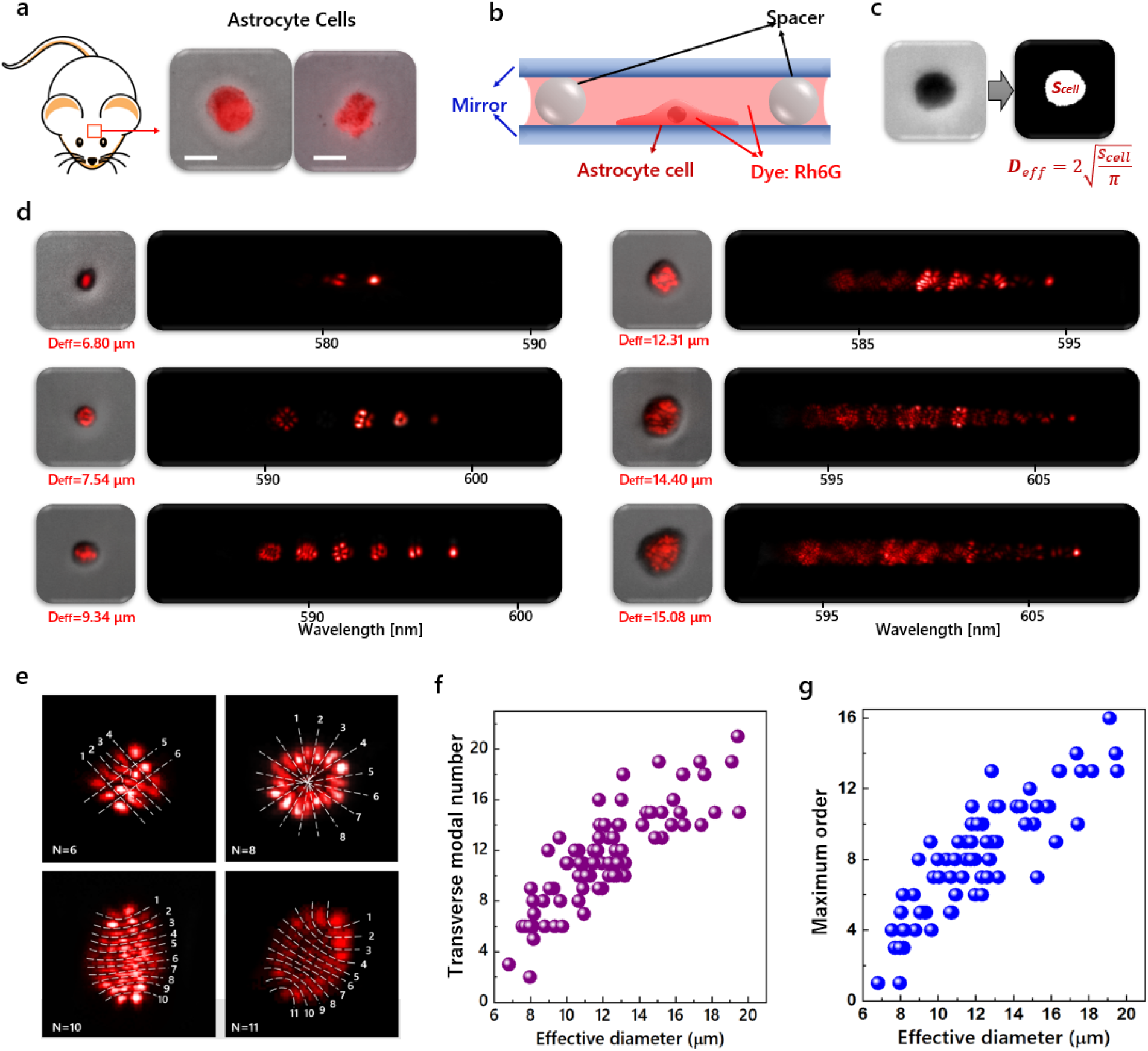
Laser modes generation from biological cells. **(a)** Fluorescent images of biological cells, which are superimposed on the bright-field images. The biological cells were fixed astrocyte cells obtained from mice nervous systems. The fluorescent dye was Rh6G. Scale bar: 20 μm **(b)** Schematic of a FP cavity sandwiching an astrocyte cell. Rh6G which was used as the lasing dye was distributed both in the cell and the surrounding environment. Microspheres with 16-μm diameters are used as spacers to fix the cavity length. **(c)** Cell size measurement. The cell area was firstly obtained by converting the bright-field image of a cell (left) into the binary image (right). As a consequence, the effective cell diameter *D*_*eff*_ was defined by2√(*S*_*cell*_/π), where *S*_*cell*_ is the cell area. **(d)** laser emission patterns superimposed on the bright-field images (left) and the corresponding hyperspectral images of laser modes (right). **(e)** Examples of laser modes from biological cells with different orders of *N*. The order *N* was defined by the total number of nodal lines in a laser mode pattern. **(f)** Measured transverse modal number of laser modes as a function of effective cell diameter. **(g)** Measured max order of laser modes (*N*) as a function of effective cell diameter.

In order to quantitively explore how cell sizes influence the output laser modes of single cell lasers, the effective cell diameter was defined to represent the cell size (Fig. 5c). The overall cell area was obtained by converting the bright-field image into a binary image. Consequently, the effective cell diameter was defined as *D*_*eff*_=2√(*S*_*cell*_/π), where *S*_*cell*_ is the cell area. By exciting the cells with 532-nm pulsed laser under a fixed pump energy density, laser modes emitted from astrocytes with different effective diameters were recorded in Fig. 5d. Similar to artificial cell models in Fig. 2c, the far-field laser emission patterns (left part in Fig. 5c) become larger and more complex as the size of astrocytes increases. Meanwhile, the corresponding hyperspectral images of laser modes (right part in Fig. 2c) also showed increasing transverse modal number of laser modes.

As shown in Fig. 5e, the laser modes cannot be well described by typical HG, LG or IG modes. This is because the irregular morphology of real cells contributes to a more complicated spatial intensity. However, we can still define the order (*N*) of laser modes by the number of nodal lines in a mode pattern (Fig. 5e). We further performed the statistics of the transverse modal number and maximum order of laser modes from astrocytes with different effective cell diameters. As shown in Figs. 5f and 2g, both the transverse modal number and the maximum order of laser modes increase with the effective cell diameters. This phenomenon is similar to the results obtained from bioinspired cell models in Figs. 2e and 2f.

### 2.6 Size prediction of biological cells from laser mode imaging enabled by deep learning

Finally, the effective diameters of astrocytes were predicted based on laser mode imaging by employing deep learning method. Herein both the far-field laser emission patterns and its corresponding maximum order laser mode component were extracted as laser mode images to train the CNN model (Fig. 6a). Note that the CNN model was also designed based on VGG19 as shown in Fig. 4b (details in Method). The CNN was trained with 1400 laser mode images in the training dataset and validated with 200 images in the validation dataset. Figure 6b plots the MSE between the measured and output effective cell diameters for training and validation datasets. Both curves show convergence to low values with increasing number of epochs. To demonstrate the performance of deep learning, we randomly selected 400 laser mode images from astrocyte cells to test the predictions of effective cell diameters delivered by the trained CNN model. The scatter plots in Fig. 6c shows the prediction results with respect to the measured effective diameters. It can be observed that the predicted points are well distributed along the 45-degree line which shows the equality between measured and predicted effective diameters. Based on the prediction data, we calculated the MAE and *R*^2^ value (details in Method), which equals 0.88 μm and 0.8471, respectively. The prediction MAE for cells demonstrated that a sub-wavelength accuracy was also achieved through the trained CNN. Figures 6d and 6e show two examples from the test dataset: the left panel represent the input laser mode images, while the right panel shows the comparison between predicted and measured cell diameters. Note that the red dashed circles in the bright field images have the same area with the astrocyte cells; and the blue dashed circles are constructed based on the predicted effective cell diameter, which are very close to the red ones. Such high coincidences between the predicted and measured effective cell diameters show excellent predictions for laser mode images made by the CNN model. The results in Fig. 6 demonstrate that a high-accuracy prediction of cell sizes could still be achieved even for biological cells with irregular shapes and random morphologies.

**Figure 6.**
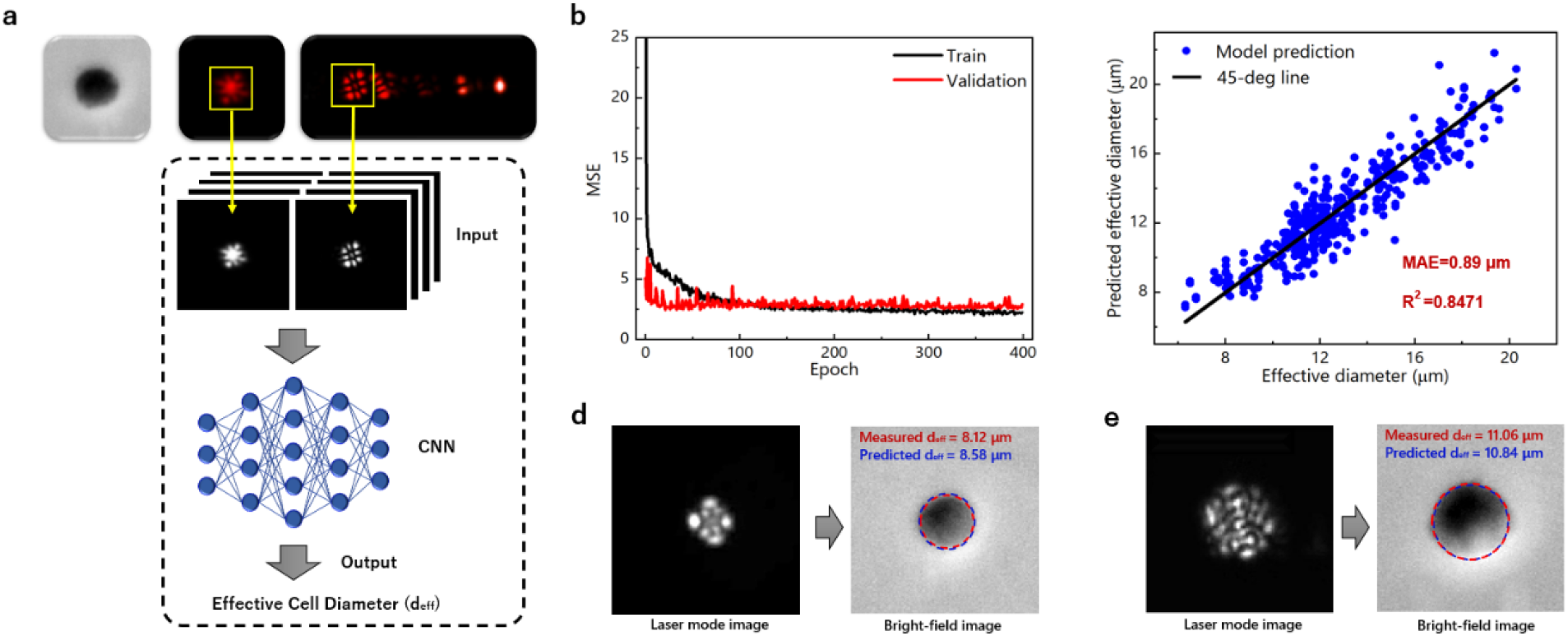
Size prediction of biological cells based on laser mode imaging using deep learning. **(a)** Schematic of size prediction of biological cells using CNN. The far-field laser emission patterns and the corresponding max-order laser mode components were selected to train a CNN model; subsequently, the effective cell diameters were predicted by the CNN model based on the laser mode images. **(b)** The MSE for training and validation sets with respect to the epochs. **(c)** Scatter plot showing test set predictions made by the trained CNN model. Blue dots: predictions of effective cell diameters. Black line: 45-degree line which represents the measured effective cell diameters. The prediction MAE was 0.89 μm; the prediction R^2^ value was 0.8471. **(d) (e)** Two examples of laser mode images (left) and the corresponding predicted effective cell diameters (right). Red dashed circles in the bright-field images have the same enclosed areas as the cells, i.e., the diameters were the measured effective cell diameters. Blue dashed circles in the bright-field cell images: reconstructed circles with predicted effective cell diameters.

## 3. Discussion and Conclusion

In this study, we explored the potential of using laser modes generated from single cell biological lasers to study critical physical features of cells. As a proof-of-concept, we demonstrated the predictions of cell sizes based on laser mode imaging by using deep learning method. Firstly, spherical cell models fabricated by CLC droplets were used to systematically study how the target sample diameters affect the emitted laser modes. Both the transverse modal number and corresponding maximum order of laser modes from FP cavities present an increasing tendency with the increasing droplet diameters. The huge diversity of laser modes based on the same droplet size was also demonstrated to show the necessity of introducing deep learning method for laser mode imaging. Consequently, laser mode images were used to train a CNN model where predictions of cell model diameters with a sub-wavelength accuracy were achieved. Finally, we demonstrated the power of laser mode imaging of biological cells by using real biological cells. Laser mode images from astrocyte cells derived from mice nervous system were used to train a CNN model. Predictions of cell sizes with a sub-wavelength accuracy were also achieved.

Our results show that subtle changes of cell physical properties (ex: cell diameter) could be amplified through FP cavities and reflected in laser mode images. Most importantly, deep learning technology has the potential to endow laser modes with biological significance and functions. The fact that cell size is linked with many important cell activities such as cell growth, proliferation, and carcinogenesis^30–33^, we envision that the established laser mode imaging system can be applied to detect various cell activities powered by deep learning. Due to whole-body interactions between cells and electromagnetic field in FP cavities, fundamental correlations between laser modes and other cell properties such as cell shape, 3D morphology, RI distribution, and cell internal spatial structures can also be built by employing deep learning. For example, the 3D morphology of a cell illustrates the boundary conditions in FP cavities, which determines the eigen solutions of transverse oscillation modes; therefore, laser modes may be applied to reconstruct 3D cell morphologies by using CNN to learn the underlying physics hidden in laser modes. In addition to biophysical properties of cells, the same concept can be used to study biochemical functions within cell. For instance, by labeling the cell nucleus, the same concept can be used to study the laser modes from size changes of cell nuclei. By labeling the cytoskeleton in cell, one can study the morphological or mechanical changes of cells through laser modes. Overall, this work illustrates the great potential of laser mode imaging of cells, offering a novel way for applications in single cell analysis and biophysical studies.

## 4. Materials and Methods

### 4.1 Materials

Nematic liquid crystal (4′-pentyl-4-biphenylcarbonitrile, 5CB), chiral dopant [4-cyano-4’-(2-methylbutyl) biphenyl, CB15] and DCM were purchased from Tokyo Chemicals. polyvinyl alcohol (PVA), Rh6G, Poly-d-Lysine (PDL) (P0899), DNase 1 type IV and Triton X‐100 were purchased from Sigma ‐ Aldrich. Alexa-Fluor 488 goat anti-rabbit, DAPI (4′,6‐diamidino‐2‐ phenylindole), paraformaldehyde (PFA, 7230681), goat serum, Dulbecco's Modified Eagle Medium (DMEM), penicillin/streptomycin (Pen/Strep) and fetal bovine serum (FBS) were purchased from Life Technologies. Rabbit anti‐GFAP (Z0334) was obtained from DAKO. Papain suspension was purchased from Worthington. Cell strainer (70 μm) was obtained from BD, Biosciences, USA.

### 4.2 preparation of CLC droplets (Cell model)

Nematic liquid crystal, chiral dopant and DCM dye were mixed and immersed in an ultrasonic bath for 15 min to form a homogenous CLC mixture with 25 wt% of chiral dopant and 0.1 wt% of DCM dye. A surfactant solution was then prepared by adding 1wt% PVA in DI water for providing surface anchoring in the formation of CLC droplets. CLC droplets were finally obtained after adding the CLC mixture in the surfactant solution with the volume ratio of 1:9 and shaking for 3 min by a vortex mixer.

### 4.3 preparation of astrocyte cells

The animal care and experimental procedures were carried out in accordance with the Institutional Animal Care and Use Committee guidelines of Nanyang Technological University (IACUC, NTU, Protocol number A19004). Briefly, P0-P2 neonatal rat cortices were isolated and digested with 1.2 U papain and 40 μg/mL of DNase at 37 oC for 1 hour. Thereafter, the digestion process was stopped by adding in 8 mL of DMEM with 10% FBS. The dissociated tissues were triturated with a syringe connected with a 21G needle, followed by a 23G needle for 10 times each. Tissues from 6 digested cortices were seeded on four PDL-coated (100 μg/mL) T75 flasks and cultured in DMEM with 10% FBS and 1% Pen/Strep for further separation of glial cells. The medium was changed every three days.

After 9-11 days of culture, the four T75 flask were placed on an orbital shaker and shaken (200 rpm, 37 °C) for 17-19 hours to remove the microglial and oligodendrocyte progenitor cells (OPCs). The remaining cells on the flasks were trypsinzed with 2 mL of 0.25% trypsin-EDTA at 37 oC for 5 min to obtain the astrocytes. The enzymatic activity was stopped by adding in 8 mL of DMEM containing 10% FBS. The cells were then spun down at 300 g for 5 min. Thereafter, the astrocytes were resuspended in 3 mL of DMEM containing 10% FBS and 1% Pen/Strep and triturated with a 1 mL pipette for 10 times before they were passed through a 70 μm cell strainer. For cell purity check (Supplementary Information), the collected cells were seeded on the PDL coated (100 μg/mL) 24-well plate at a density of 50,000 cells per well and maintained in DMEM containing 10% FBS and 1% Pen/Strep for overnight. On the next day, the cells were fixed with 4% PFA for 20 min prior to subsequent immunofluorescent staining.

For the cell laser experiment, the collected astrocytes were spun down at 300 g for 5 min to discard the supernatant. The cells were then fixed with 4% PFA for 20 min before they were spun down (300 g, 5 min) and resuspended in 1 × PBS for subsequent usage. Rh6G dye was used to stain the fixed astrocyte cells. 0.48 mg Rh6G powder was dissolved by 500 μL alcohol and diluted by 500 μL DI water. Thus, the Rh6G concentration of the final solution is 2mM. The fixed astrocyte cells were finally stained by adding the Rh6G solution in the PBS solution storing fixed astrocyte cells with the volume ratio of 1:1.

### 4.4 Optical system setup

The cavity mirrors used in this work possess a high-reflectivity in the spectral band from 570 nm to 630 nm. The pump source was a pulsed ns-laser (EKSPLA PS8001DR) integrated with an optical parametric oscillator (repetition rate: 50 Hz; pulse duration: 5 ns). An upright microscopic system (Nikon Ni2) with 10X 0.3 NA objective was used to focus the pump light onto a cell model or a biological cell, and was also used to collect the emitted laser. The pump beam diameter at the objective focal plane was measured to be 40 μm. For cell models, the laser modes were excited by pumping at the centers of cell models with 480-nm pump wavelength and a fixed pump energy density of 78 μJ/mm^2^. For biological cells, the laser modes were excited by pumping at the centers of cells with 532-nm pump wavelength and a fixed pump energy density of 149 μJ/mm^2^. The collected laser emissions from a FP cavity were separated by a beam splitter and incident into a charge-coupled device camera (Newton 970 EMCCD) and imaging spectrometer (Andor Kymera 328i).

### 4.5 Simulations of laser mode generation

The laser modes generation from FP cavities sandwiching cell models were simulated with finite element method using Comsol Multiphysics software. The Eigenfrequency Study was applied in the Electromagnet Waves, Frequency Domain interface within the Wave Optics modules, in which 2D model was employed. The refractive indices of the cell models and the surrounding environment were set as1.53 and 1.33, respectively.

### 4.6 Deep learning architectures

In this study, regression tasks were performed by employing CNN, in which the sizes of artificial cell models and biological cells were predicted based on input laser mode images from FP lasers. The operation performed by the CNN can be mathematically summarized as:

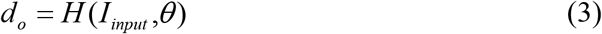

where *d*_o_ is the output value of a diameter from the CNN model; *I*_*input*_ is the spatial intensity distribution of a laser mode image which is described in Eq. 2; *θ* is the network variables; and *H* is the mapping function of the CNN model, which is composed of multiple convolutional layers, pooling layer and non-linear activation layers. In the training processes, the objective function for the regression tasks reads as:

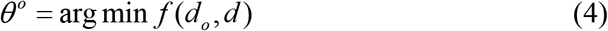

where *θ*^*o*^ is the optimized network variables; *f* is the cost function; and *d* is the labeled diameter which represents the ground truth. Equation 4 illustrates that the network variables were optimized to minimize the cost function. In this work, the mean squared error (MSE) was used as the cost function, which is defined as:

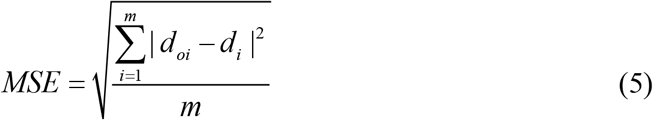

where *m* is the total number of samples; *d*_*oi*_ is the output diameter of the *i*-th sample; and *d*_*i*_ is the labeled diameter of the *i-*th sample. In this study, scholastic gradient descent (SGD) method was used to find the optimized network variables (details in Supplementary Information). For the optimization, the batch size was set to be 16, and the step size was set to be 1×10^−5^. The maximum number of epochs was fixed to be 400.

Due to limited training samples, the transfer leaning was applied in this work. The transfer learning refers to the machine learning technique that leverages learning capabilities from one task to generalize to the other new task, which is a very promising approach to help train a good network when there are limited labelled training samples. A widely used method in Computer Vision builds up the model from pre-trained network on ImageNet dataset for natural image classification task. The learned network weights are reused in the network designed for the new task, therefore reducing the number of trainable weights. Through the transfer learning, the model is able to re-use the weights from pre-trained model, yet to adapt to the new task of diameter prediction using limited number of laser mode images in this work.

The network structure was designed based on VGG19, as shown in Fig. 4b. Since the original VGG19 network was designed to classify 1000 classes of different objects from natural images, we change the fully connected layers using two new fully connected layers with smaller number of hidden nodes as shown in Fig. 4b. The weights of the newly added fully connected layers are initialized randomly. During the optimization process, the last layer of the convolutional layer and the two fully connected layers are updated while the weights in remaining layers keep the same with the original pre-trained model. VGG19 has 5 convolutional blocks, which are composed of multiple contiguous convolutional layers in the same spatial size. The max pooling operation is performed for each block to reduce the spatial size. Since in this work, the task is to predict the value of the diameter in 1-D format, the network has been modified accordingly for this task. Specially, a flatten operation is used to unfold the feature map after pool5 layer, leading to a lengthy vector. A fully connected layer (fc6) with 64 hidden nodes is added after the flatten layer and the final fully connected layer (fc7) will map the vector into the final output value to predict the diameter. A dropout layer is also added after fc6.

The prediction results were evaluated by the mean absolute error (MEA) and R^2^ value, which are defined as:

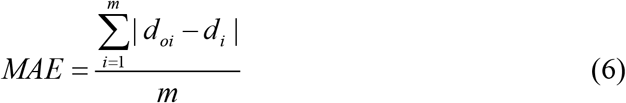

and

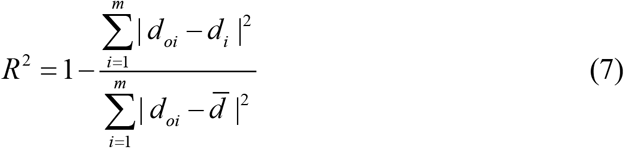

where 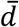 is the average diameter of the labeled samples. Note that the *R*^2^ value is between 0 and 1 and it measures how well the regression model fits the labels. The greater *R*^2^ value indicates the better the regression model fits the ground truth labels.

## Acknowledgement

This research is supported by A*STAR under its AME IRG Grant (Project No. A2084c0063). We would like to thank the lab support from Centre of Bio-Devices and Bioinformatics and Internal Grant NAP SUG from Nanyang Technological University.

## Conflict of interest

The authors declare no competing financial interests.

## Author contributions

Z. Q. and Y.-C. C. conceived the research and designed the experiments; Z. Q. and R. A. J. performed the experiments; W. S. performed the deep learning; N. Z. and S. Y. C provided the biological cells. Z. Q. and Y.-C. C analyzed data and co-wrote the paper.

## Notes

### Competing Interest Statement

The authors have declared no competing interest.

## Reference

1. Chen, Y.C. & Fan, X. Biological Lasers for Biomedical Applications. Adv. Opt. Mater. 7, 1900377 (2019).

2. Chen, Y.-C. et al. Laser-emission imaging of nuclear biomarkers for high-contrast cancer screening and immunodiagnosis. Nat. Biomed. Eng. 1, 724–735 (2017).

3. Chen, Y.-C., Chen, Q., Zhang, T., Wang, W. & Fan, X. Versatile tissue lasers based on high-Q Fabry–Pérot microcavities. Lab Chip 17, 538–548 (2017).

4. Gather, M.C. & Yun, S.H. Single-cell biological lasers. Nat. Photon. 5, 406–410 (2011).

5. Gourley, P.L. & Naviaux, R.K. Optical phenotyping of human mitochondria in a biocavity laser. Ieee Journal Of Selected Topics In Quantum Electronics 11, 818–826 (2005).

6. Martino, N. et al. Wavelength-encoded laser particles for massively-multiplexed cell tagging. Nat. Photon., 720–727 (2019).

7. Septiadi, D. et al. Biolasing from Individual Cells in a Low-Q Resonator Enables Spectral Fingerprinting. Advanced Optical Materials 8 (2020).

8. Jonas, A. et al. In vitro and in vivo biolasing of fluorescent proteins suspended in liquid microdroplet cavities. Lab Chip 14, 3093–3100 (2014).

9. Schubert, M. et al. Lasing within live cells containing intracellular optical micro-resonators for barcode-type cell tagging and tracking. Nano Lett. 15, 5647–5652 (2015).

10. Schubert, M. et al. Monitoring contractility in cardiac tissue with cellular resolution using biointegrated microlasers. Nature Photonics 14, 452-+ (2020).

11. Chen, Q. et al. An integrated microwell array platform for cell lasing analysis. Lab Chip 17, 2814–2820 (2017).

12. Schubert, M. et al. Lasing in Live Mitotic and Non-Phagocytic Cells by Efficient Delivery of Microresonators. Sci. Rep. 7, 40877 (2017).

13. Gong, C. et al. Distributed fibre optofluidic laser for chip-scale arrayed biochemical sensing. Lab Chip 18, 2741–2748 (2018).

14. Hou, M. et al. DNA melting analysis with optofluidic lasers based on Fabry-Pérot microcavity. ACS sensors 3, 1750–1755 (2018).

15. Fan, X. & Yun, S.-H. The potential of optofluidic biolasers. Nat. Methods 11, 141–147 (2014).

16. Chen, Y.-C., Chen, Q. & Fan, X. Lasing in blood. Optica 3, 809–815 (2016).

17. Humar, M. & Yun, S.H. Intracellular microlasers. Nat. Photon. 9, 572–576 (2015).

18. Van Nguyen, T. et al. Protein-based microsphere biolasers fabricated by dehydration. Soft Matter 15, 9721–9726 (2019).

19. Nizamoglu, S. et al. A Simple Approach to Biological Single-Cell Lasers Via Intracellular Dyes. Advanced Optical Materials 3, 1197–1200 (2015).

20. Humar, M., Gather, M.C. & Yun, S.-H. Cellular dye lasers: lasing thresholds and sensing in a planar resonator. Opt. Express 23, 27865–27879 (2015).

21. Moen, E. et al. Deep learning for cellular image analysis. Nature Methods 16, 1233–1246 (2019).

22. Jonsson, B.A. et al. Brain age prediction using deep learning uncovers associated sequence variants. Nature Communications 10 (2019).

23. Isozaki, A. et al. A practical guide to intelligent image-activated cell sorting. Nature Protocols 14, 2370–2415 (2019).

24. Xue, Y.J., Cheng, S.Y., Li, Y.Z. & Tian, L. Reliable deep-learning-based phase imaging with uncertainty quantification. Optica 6, 618–629 (2019).

25. Wang, Z.J., Walsh, A.J., Skala, M.C. & Gitter, A. Classifying T cell activity in autofluorescence intensity images with convolutional neural networks. Journal Of Biophotonics 13 (2020).

26. Rivenson, Y., Wu, Y.C. & Ozcan, A. Deep learning in holography and coherent imaging. Light-Science & Applications 8 (2019).

27. Jian Zhao, Y.S., Hongbo Zhu, Zheyuan Zhu, Jose E. Antonio-Lopez, Rodriog Amezcua Correa, Shuo Pang, Axel Schulzgen. Deep-learning cell imaging through Anderson localizing optical fiber. Advanced Photonics 1, 066001 (2019).

28. Rahmani, B., Loterie, D., Konstantinou, G., Psaltis, D. & Moser, C. Multimode optical fiber transmission with a deep learning network. Light-Science & Applications 7 (2018).

29. Liu, Z.W., Yan, S., Liu, H.G. & Chen, X.F. Superhigh-Resolution Recognition of Optical Vortex Modes Assisted by a Deep-Learning Method. Physical Review Letters 123 (2019).

30. Dai, X.F., Shen, Z.C., Wang, Y.H. & Zhu, M.L. Sinorhizobium meliloti, a Slow-Growing Bacterium, Exhibits Growth Rate Dependence of Cell Size under Nutrient Limitation. Msphere 3 (2018).

31. Cadart, C., Venkova, L., Recho, P., Lagomarsino, M.C. & Piel, M. The physics of cell-size regulation across timescales. Nat. Phys. 15, 993–1004 (2019).

32. Lloyd, A.C. The regulation of cell size. Cell 154, 1194–1205 (2013).

33. Maciak, S. & Michalak, P. Cell size and cancer: a new solution to Peto’s paradox? Evolutionary Applications 8, 2–8 (2015).

